# Quantifying the Fitness Benefit of Learning in Changing Environments

**DOI:** 10.1101/2022.12.16.520630

**Authors:** Emerson Arehart, Frederick R. Adler

## Abstract

The costs and benefits of learning for a foraging organism are difficult to quantify, and depend sensitively on the environment. We construct a minimal mathematical model of learning in which a forager learns the quality of different food types through experience. In our model, learning depends on two parameters: rate of memory updating and rate of exploration. Our method returns optimal learning parameters for environments in which the value and variance of food types may change in any fashion. We analyze the effect of five components of environmental change on the optimal memory and exploration parameters. The fitness outcomes from learning foragers are compared to the outcomes from following fixed strategies, explicitly quantifying the fitness benefit (or cost) of learning as a function of environmental change. We find that variance in resource values negatively biases foragers’ estimates for those values, potentially explaining experimental results showing that animals prefer less variable resources. Learning is beneficial only if memory and exploration are optimized. The benefit of learning is largely determined by the ratio between the overall expected value of taking one resource compared to the overall expected value of taking the other: As these two expectations diverge, the fitness benefit of learning decreases, and can even become negative. In many environments, sub-optimal learning performs as bad or even worse than following a fixed strategy.

**Significance:** Learning is commonly observed in foraging organisms. However, measuring the fitness benefits (and costs) of learning is difficult, and depends critically on the environment in which an organism lives. We build a minimal model of learning in the context of optimal foraging and optimal diet choice theory, with two learning parameters: *α*, corresponding to the duration of the forager’s memory, and *ϵ*, corresponding to how much the forager explores the environment to learn more about it. We identify the optimal *α*, *ϵ* for different types of environmental change, and quantify the benefits and costs of learning. The benefit of learning is often surprisingly small, and in many environments, learning provides lower fitness than following a fixed strategy.

## Introduction

Many organisms are capable of learning – acquiring information and using it to modify behavior. As with any behavioral trait, learning will persist only if it is beneficial (Dukas, 1998). However, disentangling the costs and benefits of learning is not straightforward. The value of learning depends on the environment in which it takes place. In a constant environment, learning offers no benefit; there is nothing new to learn, and any investment in learning can only be neutral or costly – for example, via increased metabolic costs associated with learning ability (Mery and Kawecki, 2003). At the other extreme, learning is of little use in a truly unpredictable environment, as there are no patterns to exploit through gathering information. In between these extremes lie environments in which learning may be beneficial (Dunlap and Stephens, 2016).

Learning is not a one-size-fits-all tool. Depending on the environment, an organism may benefit most from long- or short-term memory, or from rarely or frequently exploring the environment. A learning strategy that is optimal in one environment may be worse than not learning in another. In a rapidly changing environment, strategies which respond quickly may be optimal. In a highly variable environment, learners with a more cautious approach might fare better, because they will not constantly be chasing noise instead of signal. To analyze the fitness benefit of learning, it is therefore necessary to quantify the relationships between an organism’s fitness, how it learns, and the environment or environments in which it lives.

Optimal foraging theory is a well-developed field offering a suite of results related to optimizing strategies and the role of learning in foraging (Stephens et al., 2008). Foraging is relatively easy to manipulate experimentally (Pyke et al., 1977; Krebs et al., 1983), and associative learning is the subject of much empirical study (Dukas and Duan, 2000; Morand-Ferron, 2017). We investigate the role of learning in foraging and diet choice because it offers a straightforward metric for fitness: The total food value collected by following different strategies in the same environment. In our framework, an *environment* is specified by the reward value of two food types, how variable those rewards are, and how their values change over time. For example, the value of one food type may be held constant while another oscillates, or both types could switch stochastically among values. A foraging *strategy* is a decision rule, such as “always take one particular resource,” “select resources at random,” or “always take the resource believed to have highest value.” The strategy that results in the most food uptake over the same period of time is considered optimal.

A learning forager estimates the quality of two types of food based on experience. The forager must sample each food type to assess its quality, and decides which type of food to collect based on these estimates. Maximizing food collected is not equivalent to maximizing the accuracy of knowledge about the environment: By following the decision rule “take the resource believed to be of highest value,” the forager will behave optimally as long as it ranks the resources correctly (Arehart et al., 2022).

Two parameters determine the learning strategy for the forager: *α*, corresponding to memory, and *ϵ*, corresponding to probability of exploration. The challenge for the forager is to update the ranking of resource types as quickly as possible to reflect changes in the environment. This means that the forager must sample the environment often enough to detect changes in rank while balancing the cost of sampling against the benefit of updating. This is the “explore-exploit” tradeoff, a phenomenon of broad relevance to information gathering under uncertainty (Eliassen et al., 2007; Berger-Tal et al., 2014). Increasing precision in evaluation of resources may yield diminishing or even negative returns, as time is spent sampling non-optimal resources without providing useful information. Furthermore, updating rules must be robust to noise in the environment, and therefore able to distinguish consistent trends from random variation. A natural method for reducing variability is to average across some number of samples (the most recent 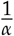 samples in our framework). A longer memory may produce more accurate estimates, but at the cost of being slow to detect persistent changes in the environment.

For each environment, we identify the pair *α*^*^, *ϵ*^*^ that maximizes food intake and the pair *α*^-^, *ϵ*^-^ that minimizes it, and compare the performance of these learning strategies with the performance of fixed strategies in the same environment. We apply this method to varying regimes of change, determining environmental conditions under which learning offers a benefit, and quantifying the benefits of learning as a function of environmental change.

Analyzing the dynamics of memory and exploration under environmental change is complex even for the most straightforward examples, and generally involves computational simulation. While simulation is a powerful tool for studying biological systems, it has important limitations. When studying dynamics with stochasticity in multiple domains – which is the case here, with variation in resource values and random sampling – many replicates must be simulated in order to identify trends. This computational intensity quickly becomes cumbersome when attempting to study global dynamics of a process. We overcome this limitation by writing probabilistic difference equations which integrate all possible learning trajectories through all possible noisy environmental conditions. While these equations cannot be solved analytically, they transform this problem into a deterministic one, enabling the use of numerical optimization to find robust solutions, and to do so at least two orders of magnitude faster than simulation.

With this work, we seek to construct a minimal model which gives insight into the fundamental mathematical structure of optimal decision-making. While this is motivated by behavioral ecology, our findings have implications well beyond animal learning. Resource allocation – of which optimal foraging is an example – is also an area of intense interest outside of biology. Our learning and decision-making framework is a special case of Temporal-Difference Learning (Sutton and Barto, 2018), as applied to a multi-armed bandit problem (Gittins, 1979) with concept drift (changing target values to be learned). Our findings could inform and increase the efficiency of related machine learning and engineering approaches.

## Results

Using the mathematical innovations developed in *Methods*, we explore environments with different regimes of change. We identify the learning parameters that maximize and minimize fitness (*α*^*^, *ϵ*^*^ and *α*^-^, *ϵ*^-^ respectively) for each environment, and compare the fitness from learning strategies to fixed strategies.

Each resource is assumed to follow a normal distribution, but the mean and variance of these distributions may change in any way over time. In the examples presented here, mean resource values *ū*, 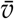 vary periodically between maxima and minima, *ū_lo_*, *ū_hi_* and 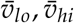 respectively, and have fixed variances *σ_u_*, *σ_v_* (Figure 1a). Learning foragers start each foraging bout without knowing the rank order of the resources, 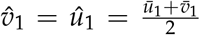. We consider two simple cases: For environments in which the timescales of change of *u* and *v* are similar, we set the period or “cycle” of *u* and *v* equal. For environments which change on very different timescales, we set *ū_lo_* = *ū_hi_*, holding *ū* constant. These cases capture the dynamics of more complex environments while being more straightforward to analyze.

**Figure 1:**
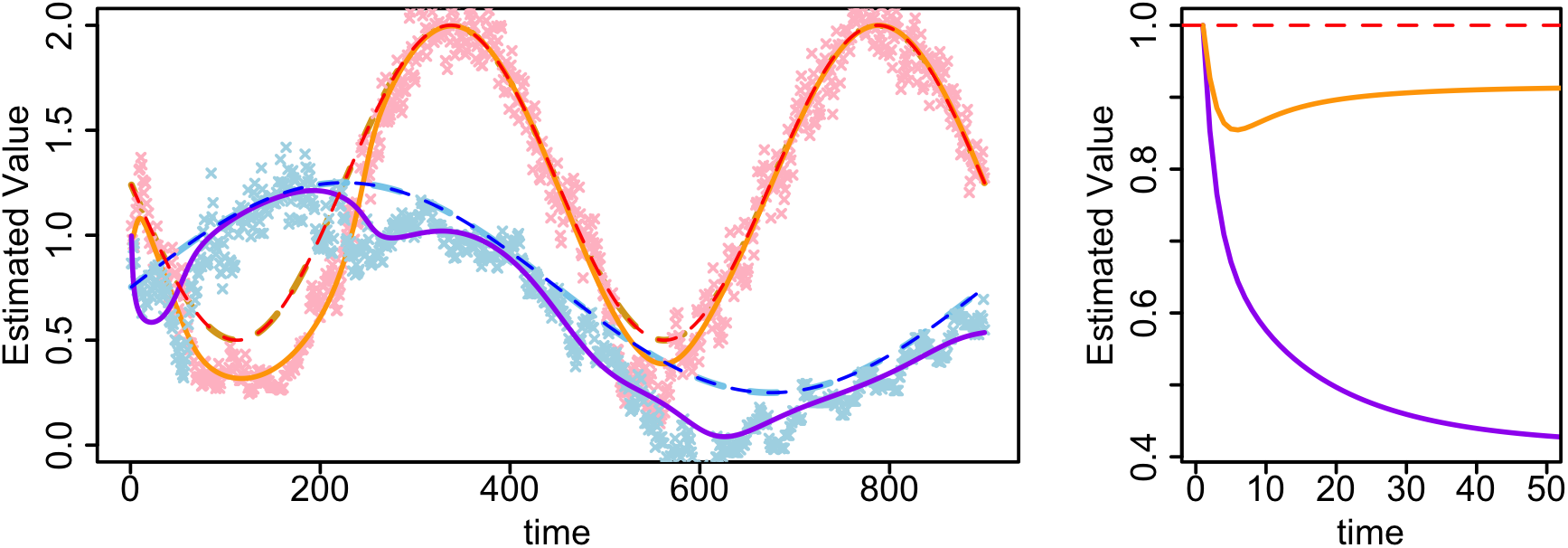
**(a)** Estimated resource values compared to true resource values over time. The blue and red dashed lines represent expected value *ū*, 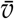 changing over time. Light blue and pink crosses show the values of *û*, 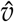 over time as obtained from multiple simulations of the foraging system. Solid orange and purple lines show *û*, 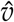 as obtained from our recursive algorithm. The forager tends to underestimate the true resource values, and the estimate *û* is distorted by 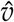. For this figure, *σ_u_* = *σ_v_* = 0.5. **(b)** Biased Sampling results in underestimation of resource values. For two resources with identical constant expected values 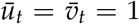 but different variance *σ_v_* = 1, *σ_u_* = 0.5, the estimate of the higher variance resource is lower. This reflects the asymmetry in sampling due to the forager’s preference for sampling higher valued resources. The rapid drop in *ū* followed by a slight ascent is not a computational artifact, but rather a result of the dynamics and interplay between the two estimates. For this figure, *α* = 0.5 and *ϵ* = 0.05.

### Dynamics of Learning

Our learning algorithm shows intriguing dynamics even with *ū*, 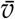 fixed. Intuitively, one might expect that the estimates *û*, 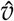 would converge on the true means *ū*, 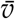 over time. However, this is not the case unless there is no variance in the resource values or 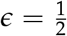, which corresponds to random sampling (which is not a learning strategy). Due to the bias induced by *ϵ*, the estimates tend to converge on values below the true expectation. The degree to which 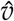 diverges from true *v* is also affected by *α, σ_v_, σ_u_*, and *û*. This results in complex dynamics, especially when both resources are changing at the same time (Figure 1).

To emphasize this, we simulate an environment where *ū* and 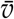 are constant and equal to 1 for all *t*, but where the variance for *v* is higher (Figure 1b). Both estimates tend towards values below the true expectation of 1, with the higher variance resource converging on a lower estimated value. This suggests a possible mechanism behind experimental findings in which foragers prefer resources with lower variance (Real et al., 1982; Cartar, 1991; Shafir et al., 1999).

### Finding Optimal α, ϵ

For each environment, we identify the combination of *α* ∈ (0,1] and 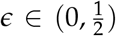 that maximizes fitness. In Figure 2, we use numerical optimization to identify optimal *α*^*^, *ϵ*^*^ pairs and plot them against change in environmental parameters: The period of cyclic change of the two resource means, *τ*, the amplitude or “span” between 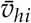 and 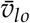 (which, in these examples, also equals the distance between *ū_hi_* and *ū_lo_*), the variance of the resource values, *σ*, and the fraction by which the *ū* cycle is offset in time compared to the 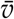 cycle. For this figure, we vary *ū* and 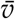 at the same rate with identical and constant variances and spans; the patterns observed here are qualitatively similar to environments where *ū* is held constant and 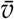 changes cyclically.

**Figure 2:**
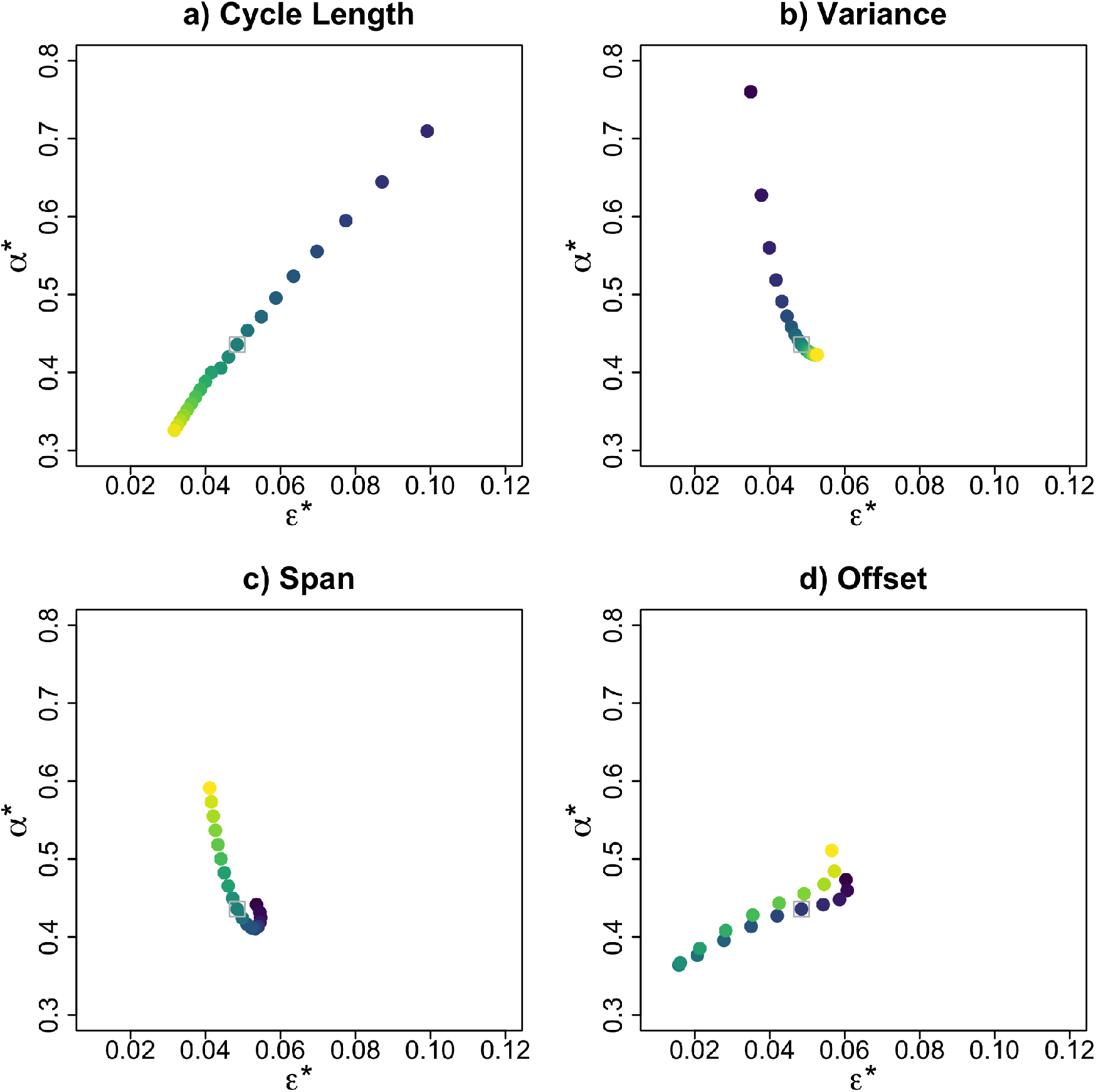
*α*^*^, *ϵ*^*^ plotted against four environmental change parameters; as values increase, dots become lighter colored (more yellow). For **(a)**, cycle length ranges from 14 to 98 timesteps in 2-step increments; in other panels, cycle length equals 100. For **(b)**, span ranges from 0.1 to 1.9 in increments of 0.1; in other panels, span equals 1. For **(c)**, variance ranges from 0.1 to 2.0 in increments of 0.1; in other panels, variance equals 1. Finally, for **(d)**, offset ranges from 0.1 to 1.9 multiplied by the cycle length, with increments of 0.1; for other panels, offset equals 0.5 times the cycle length, or halfway between perfectly in sync and perfectly out of sync.

Optimal learning depends on complex interactions between environmental parameters. However, some trends emerge from our simplified cases. In environments that oscillate quickly (small *τ*), a forager is not able to capitalize on changes in the environment, and the oscillations are essentially indistinguishable from noise. As *τ* increases, the forager is able to exploit the pattern of environmental change (Figure 2a). This manifests as a transition from learning regimes which are optimized for noisy environments to those optimized for more structured ones. As *τ* increases, *α*^*^ and *ϵ*^*^ both decrease, reflecting the more gradual regime of change and trading off the need to track the environment against noise. As variance increases, *α*^*^ decreases, reflecting the benefit of a longer memory horizon to accommodate higher noise; meanwhile, *ϵ*^*^ tends to increase, possibly because more exploration is needed to track environmental change under higher noise (Figure 2b).

The span shows a more complex trend. Initially, as span increases, *α*^*^, *ϵ*^*^ transitions from shorter memory (large *α*) and low exploration to longer memory and higher exploration, but then hooks back to shorter memory and lower exploration at higher spans. This is likely because of a phase transition in the effect of span: For very low spans, the system is driven by variance much more than by the periodic variation in 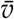. As periodic structure emerges, the value of active learning - longer memory and higher exploration - increases. At higher span, the system changes at such magnitudes that fewer samples (lower exploration) and shorter memory once again become optimal (Figure 2c).

Finally, we consider the temporal offset between the oscillations in *ū* and 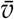, where an offset of 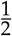 sets the resources perfectly out of phase, and offset equal to 0 or 1 means the resources are perfectly synchronized. If the two resources are perfectly synchronized, learning trivially provides no benefit (not shown). When they are slightly out of sync, *α*^*^ and *ϵ*^*^ are both relatively high, perhaps because exploration and shorter-term memory are necessary to capitalize on the small differences in resource values. *α*^*^, *ϵ*^*^ are both minimized when the oscillations are perfectly offset, as exploration is not necessary to detect changes in rank order, especially with a longer memory horizon, which will keep the estimates 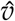, *û* close to each other (Figure 2d). For environments where *ū* is constant, the temporal offset is undefined.

### Fitness Benefits of Learning

In many environments, learning offers a fitness benefit, though the margins can be surprisingly small. Even with no costs for implementation of learning, there are environments where following a fixed strategy is as beneficial or more beneficial than even the most optimal learning. This is trivially true if the rank order of *ū_t_*, 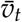 is the same for all *t*. In such cases, the fixed strategy of accepting the higher valued resource type is the theoretical maximum; learning may be able to match that, but cannot exceed it, and could in fact be pathological, for example by mistaking variation for trends, resulting in worse fitness outcomes.

The most important factor affecting the fitness benefit of learning is the ratio between the expectation of always taking *v* to the expectation of always taking *u*. As this ratio of the expected value of the fixed strategies, *E*[*v*]/*E*[*u*], diverges from 1, the margins for potential benefits from learning decrease dramatically. Not only do *α*^*^, *ϵ*^*^ sometimes yield fitness below the best fixed strategy, but the worst learning strategies *α*^-^, *ϵ*^-^ *never* exceed the best fixed strategy, often performing worse than fixed strategies by a significant margin (Figure 3). This is true for environments with fixed *ū* and environments with both resources oscillating, and underscores the importance of learning “correctly” for a given regime of environmental change. While *E*[*v*]/*E*[*u*] drives the fitness value of learning, it has only a very minor effect on determining the optimal learning parameters (and thus is not pictured in Figure 2).

**Figure 3:**
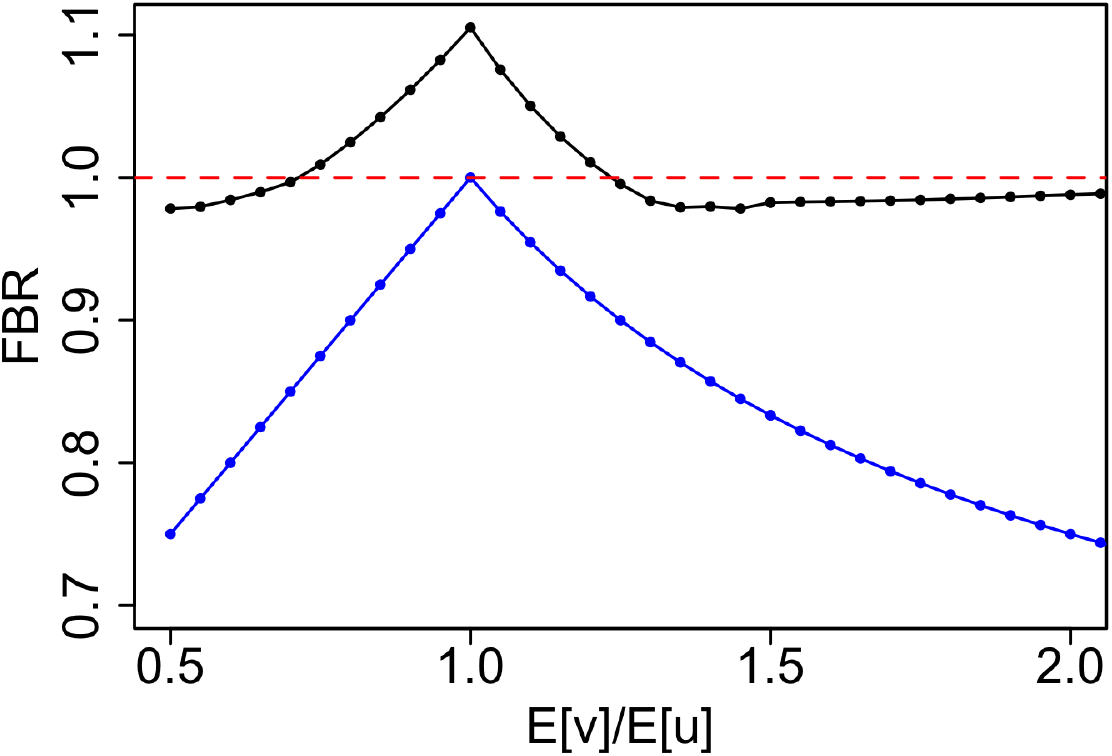
Fitness benefit ratio (FBR) plotted against the ratio between always taking *v* to always taking u, *E*[*v*]/*E*[*u*]. Black points correspond to the ratio of fitness from the best learning strategy, parameterized by *α*^*^, *ϵ*^*^, to the fitness from the best fixed strategy. Blue points correspond to the ratio of fitness from the *worst* learning strategy, parameterized by *α*^−^, *ϵ*^−^, to the best fixed strategy. For points above the dashed red line, learning provides a fitness benefit; below it, learning is worse than following a fixed strategy. The fitness benefit of learning is often marginal, and the potential detriment from learning, especially with non-optimal learning parameters, is substantial. For this figure, cycle length = 100, span = 1, variance = 1, and offset = 0.5 times the cycle length (50 timesteps).

In Figure 4, we compare the fitness values attained from our best learning strategy (parameterized by *α*^*^, *ϵ*^*^) to the fitness values from the best fixed strategy in each environment, varying across the same environmental variables as before. We also plot the fitness which results from *α*^-^, *ϵ*^-^, the *worst* learning parameters in each environment. Learning provides a greater benefit as the period of oscillation *τ* increases, as the span increases, as variance decreases, and as the two oscillations are increasingly out of phase (Figure 4). While learning enables adaptation to major environmental changes, exploration is costly, and the forager must balance this cost against the penalty for missing an environmental change and following the wrong strategy for longer, both of which increase as the cycle length becomes large and the offset between resources approaches 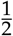. Complicating matters further, increasing the variation in the environment means that a forager must select *α* such that it captures the magnitude of change in a resource (span) but does not estimate incorrectly due to variance, again balancing the cost of updating estimates more slowly and hence being slower to switch strategies.

**Figure 4:**
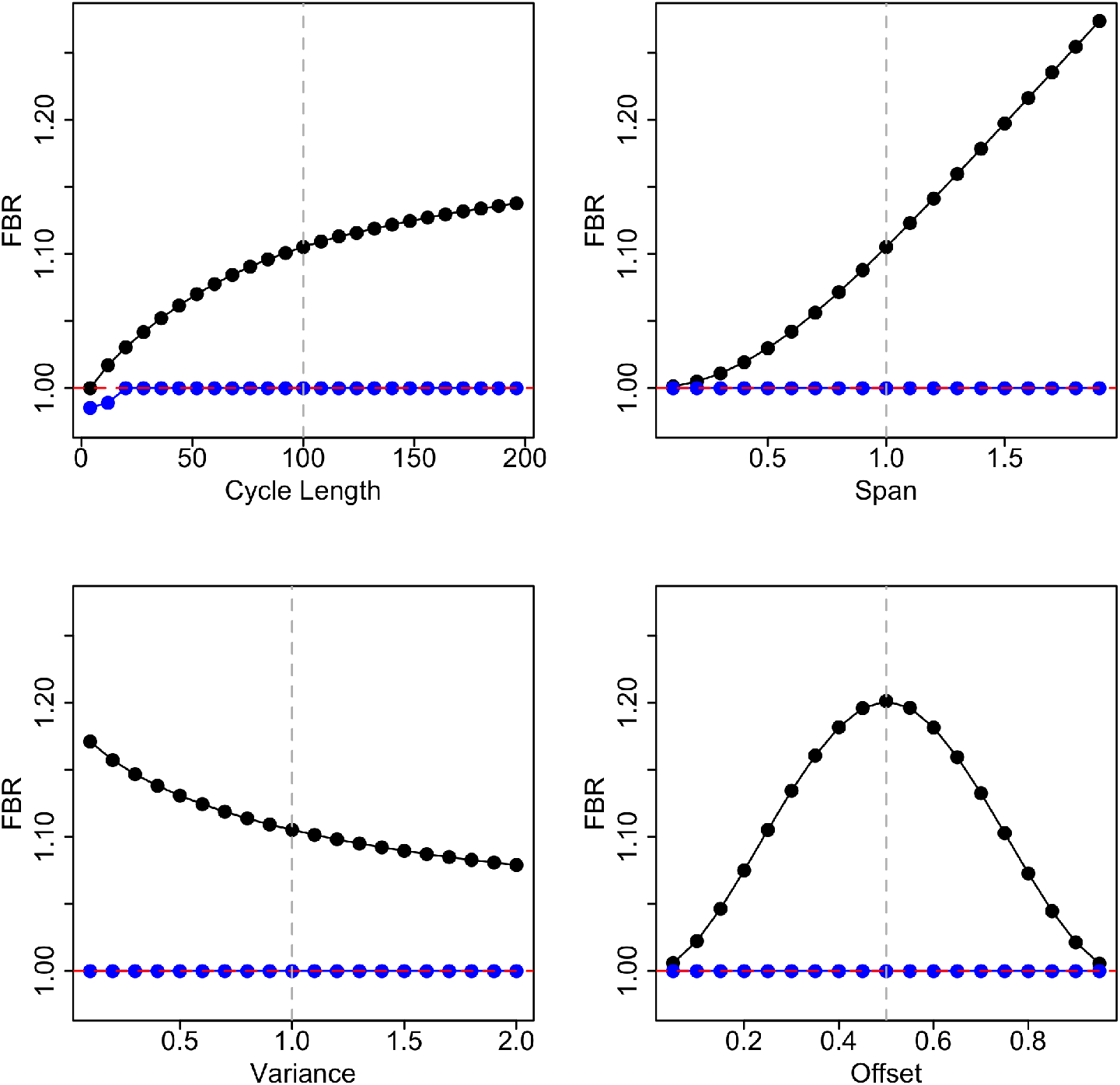
Fitness benefit ratio (FBR) plotted against environmental change parameters. Black points correspond to the ratio of fitness from the best learning strategy, parameterized by *α*^*^, *ϵ*^*^, to the fitness from the best fixed strategy. Blue points correspond to the ratio of fitness from the *worst* learning strategy, parameterized by *α*^−^, *ϵ*^−^, to the best fixed strategy. Learning provides an increasing (but saturating) benefit as cycle length increases; an increasing benefit with increasing span; decreasing benefit with greater variance; and the greatest fitness benefit when environmental change cycles are the most offset. For **(a)**, cycle length ranges from 6 to 98 timesteps in 2-step increments; in other panels, cycle length equals 100. For **(b)**, span ranges from 0.1 to 1.9 in increments of 0.1; in other panels, span equals 1. For **(c)**, variance ranges from 0.1 to 2.0 in increments of 0.1; in other panels, variance equals 1. Finally, for **(d)**, offset ranges from 0.1 to 1.9 multiplied by the cycle length, with increments of 0.1; for other panels, offset equals 0.5 times the cycle length, or halfway between perfectly in sync and perfectly out of sync.

## Discussion

A primary challenge for a learning forager is to distinguish noise from meaningful, exploitable patterns. We demonstrate that this is possible even with a minimal form of learning, in which the learner only tracks point estimates of the quality of two resources. For learning to be beneficial, it must be properly tuned to the regime of change in a forager’s environment. Learning can quickly become a liability for an organism if it is acquiring and using information sub-optimally. In many cases the best learning strategy is actually worse than the best fixed strategy. Even when learning offers a fitness benefit, the marginal benefit that learning provides over the best fixed foraging strategy is often surprisingly small.

If a forager has a fixed memory parameter *α* and a fixed learning parameter *ϵ*, perhaps optimized by evolution for a particular regime of environmental change, those parameters may be detrimental under different environmental conditions. Foragers might be able to tune *α*, *ϵ* in response to other cues from the environment; this would constitute a higher order of learning, outside of the scope of our minimal model. If foragers are unable to change *α*, *ϵ*, they may experience major fitness declines under environmental conditions for which they are not suited. This could be relevant for species experiencing ecosystem changes wrought by invasive species, habitat fragmentation and loss, and climate change. The framework presented here could be applied to test the sensitivity of a given learning parameterization to larger-scale changes in the environment. It could also be applied to obtain a “generalist” *α*, *ϵ* by optimizing learning for an environment undergoing multiple regimes of change.

We explore environmental change by simulating cyclic change at multiple scales. Our intention is not necessarily to model, for example, diurnal or seasonal change, which a forager might be able to predict indirectly from other cues, such as temperature or daylight. Rather, by simulating an environmental regime change over multiple iterations, we seek to capture the intrinsic speed and magnitude of that type of change in a robust manner.

The mathematical framework introduced here is very flexible, and can be applied to many different regimes of environmental change beyond those considered here, such as stochastic (rather than cyclic) switching of resource values. It could also be extended to more than two resource types, and additional higher-order learning parameters could be incorporated. For example, in the sampling probability equation (Equation 4), we fix the switch sensitivity parameter *k* = 10. This corresponds to smooth but confident switching, in which any significant difference in estimated resource values will result in sampling the lower-rank resource with probability *ϵ*. However, *k* could be analyzed as an additional learning parameter, where lower *k* corresponds to less confidence in a forager’s estimates, and thus increased willingness to explore.

We focused on resources which are normally distributed. However, it may be possible to extend this framework to other types of resource variability, such as resources with a binomial reward probability. The framework could also be extended to include group foraging behavior and information exchange. Here we analyzed an individual agent solving a multi-armed bandit problem with concept drift (changing reward values). Understanding how multiple individuals can optimally learn and share information – the multi-agent, multi-armed bandit problem – is an important area of study in systems and control theory (Landgren et al., 2021). By extending our framework to include information exchange between agents, it might be possible to make progress in the area of multi-agent, multi-armed bandits with concept drift.

The concepts here could be tested using experimental techniques that have been in use by optimal foraging researchers for decades. The dynamics of even such a minimal model of learning can be complex and counterintuitive, as shown in the dynamics of biased sampling on estimates, and in the ubiquity of environments in which learning may be detrimental. Researchers testing for learning must be cautious and consider such idiosyncrasies when planning experiments. Another challenge is choosing the timescale and the way that variance is implemented such that more clever organisms don’t learn the temporal structure of the switching and exploit that, as they have been shown to do in, for example, Stephens (1987). One possibility is for an organism to face a series of binary foraging decisions in the form of a y-maze (Dunlap et al., 2017). We omitted handling time and switching costs (for example, time moving between a patch of *u* and a patch of *v*) from our model for mathematical simplicity, but the general dynamics presented here are likely to still hold, albeit with different learning parameters.

Understanding how learning is actually implemented in a forager’s brain is a formidable challenge. However, we hope that this minimal model provides a beginning mathematical approach to learning that will guide future theoretical and experimental inquiry.

## Methods

We consider a forager with access to two resource (food) types, *u* and *v*. Each resource type is always available, and the value of each resource at time *t* follows an independent normal distribution with means *ū_t_*, 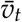 and variances *σ_ut_, σ_vt_* respectively. The parameters of these distributions can change over time in any fashion.

The total fitness from a foraging bout is the sum of the resources acquired at each time step during the bout. The expected total fitness *F* is obtained by multiplying the probability of taking resource *v* at time *t* by the expected value at that time, 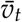, plus the probability of taking resource *u* at *t* multiplied by *ū*. For fixed strategies and total foraging time *T*, we have

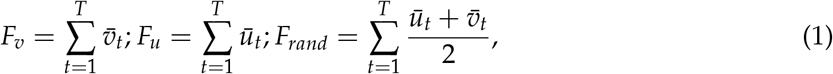

where *F_v_* is the total fitness from only taking *v*, *F_u_* is the total fitness from only taking *u*, and *F_rand_* is the total fitness from randomly choosing *u* or *v* at each time step. For comparison, we also calculate the best (*F*^*^) and worst (*F_worst_*) fitness outcomes theoretically possible in the environment

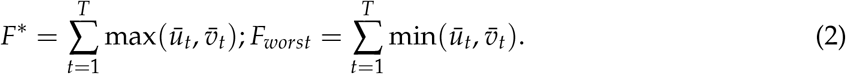

Learning foragers maintain internal estimates of the two resources, *û* and 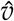; these estimates are updated only when the forager samples the corresponding resource, so only one of these quantities can be updated at any time step. The estimates are updated following a proportional update rule

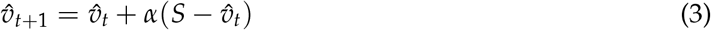

and likewise for *û*. This expression is based on the Rescorla-Wagner rule, an updating rule commonly used in studies of associative learning (Rescorla, 1972), as well as in optimal foraging theory, including McNamara and Houston (1987) and Eliassen et al. (2007) (note that *α* has the opposite interpretation in McNamara and Houston (1987)). *S* is a sample of the resource drawn from a normal distribution with mean 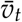 and variance *σ_vt_*. *α* is the memory parameter; smaller *α* corresponds to a longer memory, integrating information over roughly the past 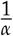 samples of the resource. *α* = 1 corresponds to the shortest possible memory, with the estimated value of a resource being exactly equal to the most recent sampled value. *α* is constrained to take a minimum value of 0 (no learning) and a maximum value of 1.

At each time step, the forager samples resource *v* with probability *p*, and always takes one resource or the other, so takes resource *ū* with probability 1 – *p*. *p* is a function of the value of the estimates *ū*, 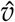. The forager wishes to maximize fitness, so will generally prefer to take the resource it estimates to be of higher quality. However, in order to gather information about the current state of the environment, the forager may sample the resource it estimates to be non-optimal. We consider two methods for the forager to determine the probability of sampling. First, in the “switch” case, the forager may explore with a fixed probability *ϵ*; if 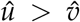, the forager will sample resource *v* with probability *p* = *ϵ* and resource *u* with probability 1 – *p* = 1 – *ϵ*. *ϵ* is constrained to take a value between 0 (no exploration) and 0.5 (randomly sampling both resources). Even if *ϵ* = 0, if *α* > 0, a forager’s strategy may still change under some circumstances. For example, if resource *v* is preferred, but the value drops such that 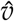 drops below *û*, the strategy will switch. The forager only follows a completely fixed strategy if *α* = 0.

For increased biological realism, we also consider the “sigmoid” case, in which the probability of sampling *v* follows a sigmoid function, meaning that if the estimated resource values are close, the probability of exploration increases. The probability of sampling *v* is given by

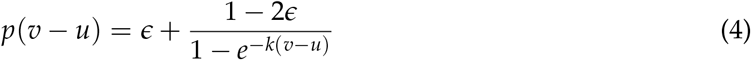

where *k* is a parameter controlling the sharpness of the cutoff; for high *k*, this approaches fixed switching. In this paper, we fix *k* = 10, but *k* could be considered an additional learning parameter.

Fitness is assessed by computing the expected value of total resources collected over a foraging bout. We compute the fitness for an environment across the range of possible *α, ϵ* combinations, and then identify the optimal *α*, *ϵ* and compare it to the fitness achieved by fixed strategies.

To compute the expected fitness in a changing environment while incorporating the variation in resource values, we implement a recursive algorithm in random variables. We outline that algorithm for *v*, but the process for *u* is symmetrical.

Denote the estimate 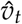 by the random variable 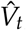, and the variance of the distribution of estimates at time *t* by the random variable *W_t_*. The value of 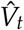 is given by the recursion

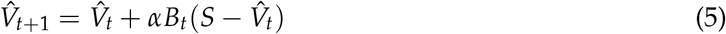

where 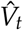 is the estimate of the value of resource *v* at time *t, S* is a random variable sampled from the normal distribution with mean 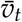 and variance *W_t_*, and *B_t_* is a Bernoulli random variable which takes on value 1 if *v* is being sampled at time *t* (which depends on *ϵ*), and value 0 otherwise. We wish to find the expected value 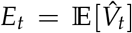 for every time step. We first do this for fixed exploration probability (“switching”) and then extend to the case with increased probability when *û_t_* and 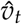 are close (“sigmoid”).

### Switching Case

Set 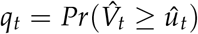. Define a Bernoulli random variable

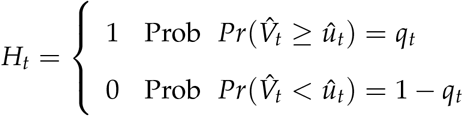

Define another Bernoulli random variable *B_t_* which will equal 1 if resource *v* is being sampled at time *t*, and 0 otherwise:

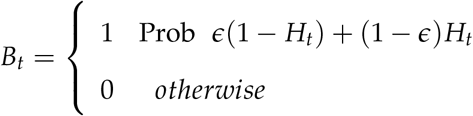

Equation (5) becomes

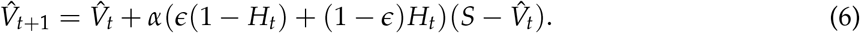

Taking the expectation of equation (6), and recalling that 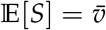, we have

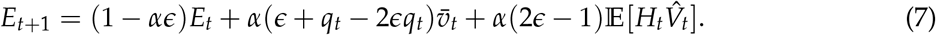

Note that 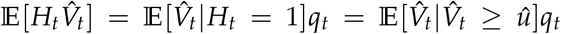. For a normal distribution, the latter quantity can be computed with the Inverse Mills ratio (Mills, 1926). Define

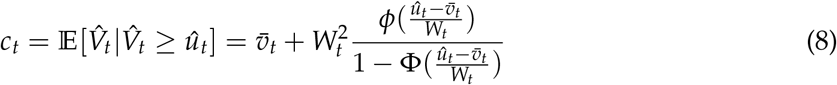

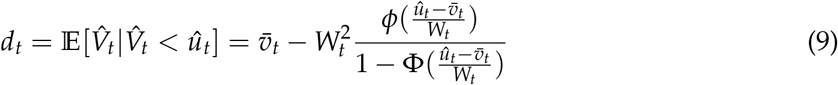

and

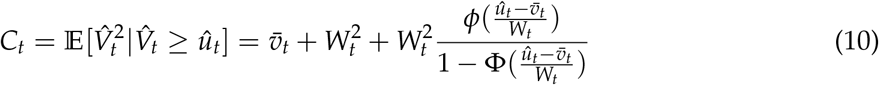

where *ϕ*(*x*) is the probability density and Φ(*x*) the cumulative distribution of a standard normal distribution. Our expression for *E_t_* then becomes

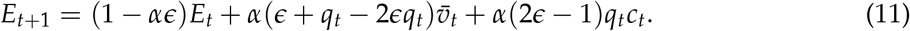

We find the variance of 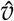 at time *t*, *W_t_*, by applying the relation 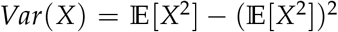 to equation (11):

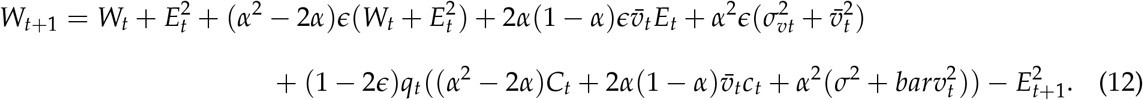

Finally, observe that *E_t_* = *q_t_c_t_* + (1 – *q_t_*)*d_t_* by definition, and solve to find *q*_*t*+1_ from *E*_*t*+1_ and *W*_*t*+1_

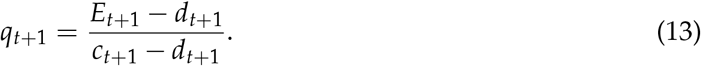

The expected fitness accrued by a forager at *t*, *f_t_*, is given by the simple relation

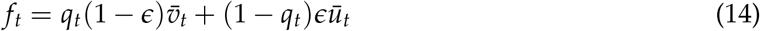

and by summing across *t*, the total expected fitness for a foraging bout is calculated.

### Sigmoid Case

To derive the equations for sigmoid exploration, we first divide the real line into two intervals, where *q_i_* represents the probability that 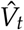 lies in interval *i* at time *t, p_i_* the probability of updating at *t* + 1 if 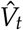 lies in interval *i*, and *c_i_* the expected value of 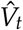 at time *t* conditional on it lying in interval *i*:

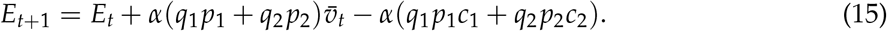

This formulation is equivalent to (11) if *p*_1_ = *ϵ*, *p*_2_ = 1 – *ϵ*, *q*_1_ = 1 – *q*, and *q*_2_ = *q*. Next, generalize to a set *I* consisting of multiple intervals:

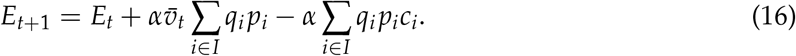

Assuming that *û* adopts a fixed value *y*, and taking the limit as the number of regions in *I* goes to infinity, we have

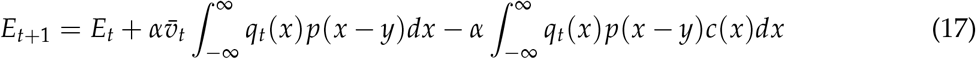

where *q*(*x*) is the probability density of 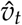, *p*(*x* – *y*) is the probability of sampling *v* if 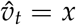 and *û_t_* = *y*. Taking the limit as the number of regions in *I* goes to infinity, 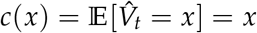, so

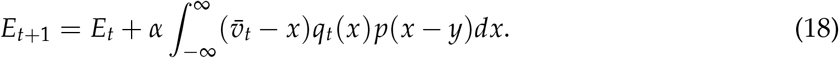

Finally, to account for the distribution of values of *u_t_*, we must integrate over the probability density *g_t_*(*y*) of *û_t_* to obtain

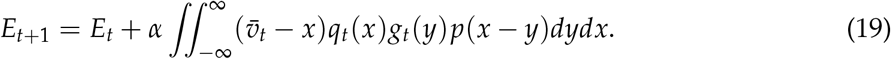

### Integral Approximation

This double integral cannot be solved analytically. However, we can reduce it to a single integral using an approximation. We rewrite the double integral as

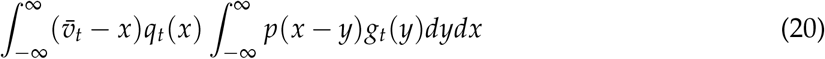

then integrate the second integral by parts (and name it *J*) to obtain

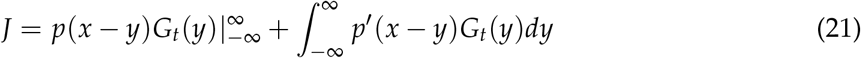

where *G_t_*(*y*) is the antiderivative of *g_t_*(*y*), which is the cumulative density of *û_t_*. The first term equals *ϵ*; for the remaining integral, we approximate the derivative of *p* with *p′* = (1 – 2*ε*)*δ*(*x* – *y*) because the sigmoid function *p* is constant at most points, adopting value *ϵ* or 1 – *ϵ*, except where *x* and *y* are close. We then have *J* ≈ *ϵ* + (1 – 2*ϵ*)*G_t_*(*x*). Following LaPlace’s Method (Bender and Orszag, 2013), we add a term to the expansion where *p′*(0) is steep. We use change of variables to rewrite *p′* as two exponential functions, 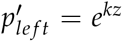 and 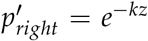, with 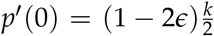, so that

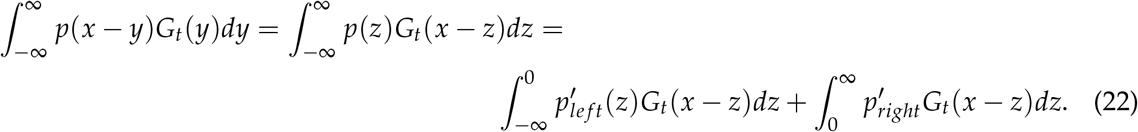

By LaPlace’s method, the farthest right integral is approximated by

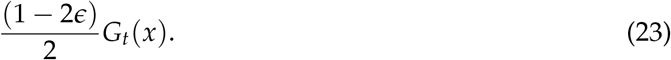

By Taylor expansion of *G_t_* near 0, we have 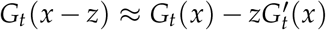. Following this procedure,

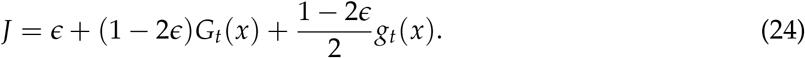

Finally, plugging this into (19), we have

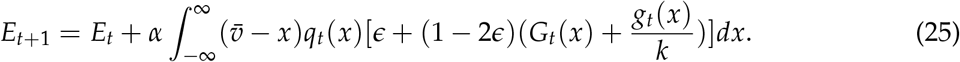

Omitting 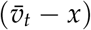 from this integral gives the total probability of sampling resource *v* at time *t*, which we call *Q_t_*:

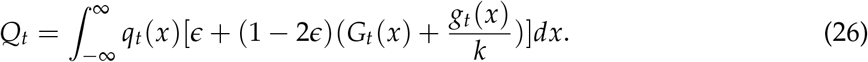

By a similar process, we can find the variance *W_t_*:

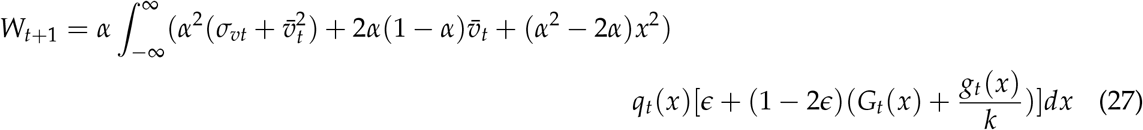

which depends on *W_t_* and *W_t_* via *q_t_* and *g_t_*. The expression for the variance of *u* is symmetric. Applying these equations recursively throughout a foraging bout, we can track the expectation and variance in estimates of resource quality *û*, 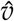, and from those extract the expected probability that each resource will be taken by a forager at each time step, *Q_t_*. To compute total fitness for pair *α*, *ϵ* in an environment defined by time series of expected values *ū_t_*, 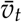 and variances *σ_ut_, σ_vt_* we obtain the probabilities of each resource being taken at every time step, and multiply by the expected value of that resource at each timestep,

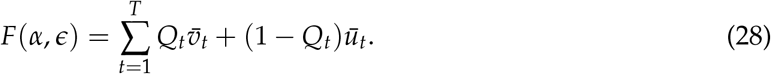

### Recursive Experiments

To determine if learning offers a fitness benefit over fixed strategies in a given environment, we first maximize *F*(*α, ϵ*) and compare it to fixed strategies *F_v_*, *F_u_*, *F_rand_*. In R (R Core Team, 2020), we solve equations (25 - 27) at each time step using numerical integration, and then use Nelder-Mead optimization to maximize *F*(*α*, *ϵ*). While we are unable to solve analytically for the optimal *α*, *ϵ*, our method allows for deterministic optimization which incorporates the natural variation in the system, and is at least two orders of magnitude faster than simulating the system with similar accuracy.

## Notes

### Competing Interest Statement

The authors have declared no competing interest.

